# Oxford Nanopore MinION genome sequencer: performance characteristics, optimised analysis workflow, phylogenetic analysis and prediction of antimicrobial resistance in *Neisseria gonorrhoeae*

**DOI:** 10.1101/349316

**Authors:** Daniel Golparian, Valentina Donà, Leonor Sánchez-Busó, Sunniva Foerster, Simon R. Harris, Andrea Endimiani, Nicola Low, Magnus Unemo

## Abstract

Antimicrobial resistant (AMR) *Neisseria gonorrhoeae* strains are common and compromise gonorrhoea treatment internationally. Rapid identification and characterisation of AMR gonococcal strains could ensure appropriate and even personalised treatment, and support identification and investigation of gonorrhoea outbreaks in nearly real-time. Whole-genome sequencing is ideal for investigation of the emergence and dissemination of AMR determinants that predict AMR in the gonococcal population and spread of AMR strains in the human population. The novel, rapid and revolutionary long-read sequencer MinION is a small hand-held device that can generate bacterial genomes within one day. However, the accuracy of MinION reads has been suboptimal for many objectives and the MinION has not been evaluated for gonococci. In this first MinION study for gonococci, we show that MinION-derived sequences analysed with existing open-access, web-based sequence analysis tools are not sufficiently accurate to identify key gonococcal AMR determinants. Nevertheless, using an *in house*-developed CLC Genomics Workbench, we show that ONT-derived sequences can be used for accurate prediction of decreased susceptibility or resistance to recommended therapeutic antimicrobials. We also show that the ONT-derived sequences can be useful for rapid phylogenomic-based molecular epidemiological investigations, and, in hybrid assemblies with Illumina sequences, for producing contiguous assemblies and finished reference genomes.

## Introduction

Gonorrhoea is a frequently asymptomatic sexually transmitted infection caused by *Neisseria gonorrhoeae* (gonococcus). In 2012, the World Health Organization (WHO) estimated that there were 78 million new cases of gonorrhoea among adults worldwide^1^. Untreated gonorrhoea can lead to serious reproductive tract and (rarely) systemic complications^2–4^. The high estimated global incidence of gonorrhoea, increasing rates of reported prevalence in many more-resourced settings, and particularly the increasing levels of antimicrobial resistance (AMR) have grown to major public health concerns^1,5–8^. Resistance to ceftriaxone and azithromycin in *N. gonorrhoeae* threatens the currently recommended dual antimicrobial therapy, mainly ceftriaxone 250-500 mg intramuscularly plus azithromycin 1-2 g single oral dose, and no new antimicrobials for treatment of gonorrhoea are yet available^4,6–15^. Consequently, WHO included *N. gonorrhoeae* in its first list of AMR “priority pathogens” in February 2017^8,16^. Therefore, timely detection and surveillance of AMR gonococcal strains, their AMR determinants and their emergence and dissemination in the populations globally is crucial^5^,^6–8^,^11^,^17–19^.

Next generation sequencing (NGS) is ideal for the elucidation of the molecular determinants of AMR, their dissemination throughout the gonococcal population, and the emergence and dissemination of AMR strains in the human population, nationally and internationally^20,21^. The demands for technologies that can operate at higher speed with less hands-on time have driven innovation into new sequencing approaches. Third-generation sequencing (TGS)^22^ with benefits such as increased read length, reduction of sequencing time and reduction of sequencing bias introduced by PCR amplification steps^23^, is being used increasingly. Most current DNA sequencing methods are based on either chemical cleavage of DNA molecules^24^ or synthesis of new DNA strands. The synthesis-based methods sequence DNA strands labeled either through the primer from which it originates or on the newly incorporated bases. This is the basis of the sequencing methods used on most currently available platforms, including Illumina, Ion Torrent (Thermo Fisher Scientific), and Pacific Biosciences (PacBio) sequencers. It has also been shown that individual DNA molecules can be sequenced by monitoring their progress through various types of pores^25,26^. The benefits of this approach include potentially very long and unbiased sequence reads, because neither amplification nor chemical reactions are necessary for sequencing^27^. Oxford Nanopore Technologies (ONT) (Oxford, UK) have introduced this approach with their genome sequencing device MinION, a TGS platform that was commercialized in mid-2015^28,29^. The MinION is a single-molecule nanopore sequencer in which nanopores are embedded in a membrane placed over an electrical detection grid. As part of the library preparation protocol, proteins are bound to the DNA fragment, and the DNA-protein complex is passed through the pore as the motor protein helps to unwind the DNA, controlling the rate of single stranded nucleotides that pass through the pore one at a time^30^. DNA molecules passing through the pores create measureable alterations in the ionic current. These fluctuations (“squiggles”) are sequence dependent so the change in ionic current across the membrane can be used by a base-calling algorithm to infer the sequence of nucleotides in each molecule^27,31^. Under ideal conditions, the template DNA strand passes through the pore, followed by a hairpin adapter and finally the complementary strand. In a run where both strands are sequenced, a consensus sequence of the molecule can be produced; these consensus reads are termed two-directional reads or “2D reads” (2D ONT reads) and generally have higher accuracy than reads from only a single pass of the molecule (1D ONT reads). The MinION provides several new advantages: small size (10 × 3 × 2 cm), portability, speed, low cost, and direct connection to a laptop through a USB 3.0 interface; the library construction involves simplified methods; no amplification step is required, and data acquisition and analysis occur in real time. The MinION is promising for microbiological applications including describing the microbiome^32^, rapid diagnostics^33^, transmission and surveillance^34,35^, *de novo* assemblies^36–38^, and microbial AMR profiling^39,40^. However, the high error rate for the MinION sequencer^28,29^ has limited its ability to compete with existing sequencing technologies and the MinION has not been previously evaluated for gonococci.

We evaluated the performance characteristics, ideal sequence analysis (tools and workflow for taxonomy, assembly, assembly improvement (“polishing”) and mapping), phylogenomic analysis, and prediction of decreased susceptibility or resistance to recommended therapeutic antimicrobials in *N. gonorrhoeae* using the Oxford Nanopore MinION genome sequencer. We sequenced the 2016 WHO gonococcal reference genomes/strains^41^ and clinical gonococcal isolates, and evaluated the performance of ONT assemblies and hybrid assemblies including both ONT and Illumina reads. The assemblies were evaluated against the PacBio RS II-sequenced 2016 WHO gonococcal reference genomes (Bioproject PRJEB14020)^41^ for sequence variation and to assess whether the assemblies were of sufficient quality for characterisation of AMR determinants and phylogenomic-based molecular epidemiology of gonococci.

## Methods

### *N. gonorrhoeae* isolates, culture, antimicrobial susceptibility testing and DNA isolation

Twenty-eight gonococcal isolates, including the previously PacBio-sequenced 2016 WHO reference strains (n=14)^41^ and 14 selected clinical AMR isolates, were examined. The clinical isolates were collected as part of the Rapid Diagnosis of Antimicrobial Resistance in Gonorrhoea (RaDAR-Go) project from patients attending clinics in Zurich and Bern, Switzerland, in 2015-2016^42^. All isolates were cultured on GC agar plates (3.6% Difco GC Media agar base (Becton Dickinson, Franklin Lakes, USA) with 1% IsoVitalex (bioMérieux, Marcy-l’Étoile, France), 1% haemoglobin (Becton Dickinson) and 10% horse serum (SVA, Uppsala, Sweden)) and incubated at 36°C, 5% CO_2_-enriched atmosphere for 18-20 h. We determined minimum inhibitory concentrations (MICs) of ceftriaxone, cefixime, azithromycin, spectinomycin, ciprofloxacin, tetracycline and benzylpenicillin using the Etest (bioMérieux), according to manufacturer’s instructions. Resistance breakpoints from the European Committee on Antimicrobial Susceptibility Testing (EUCAST; www.eucast.org/clinical_breakpoints/) were applied. Nitrocefin test (Thermo Fisher Scientific, Wilmington, DE, USA) was used to detect β-lactamase production. Genomic DNA was isolated using the Wizard Genomic DNA Purification Kit (Promega Corporation, Madison, WI, USA). Isolated DNA was quality controlled using fluorometric quantification (Qubit; Thermo Fisher Scientific) and electrophoresis (Tapestation; Agilent, Santa Clara, CA, USA).

### Oxford Nanopore Technologies library preparation and MinION sequencing

Three μg of genomic DNA was sheared to an average fragment length of 8 kb with g-TUBES (Covaris, Woburn, WA, USA). Sequencing libraries were prepared according to the 2D library preparation protocol with the SQK-LSK208 2D ligation kit (Oxford Nanopore Technologies), including the DNA repair step with the NEBNext FFPE DNA repair module (New England Biolabs, Ipswich, MA, USA). The sequencing libraries were purified using MyOne C1 beads (Thermo Fisher Scientific), and 6 μl of sequencing library were loaded onto a R9.4 SpotON flow cell and sequenced with the MinION Mk 1B sequencing device (Oxford Nanopore Technologies) for 24 hours, including a top-up with additional 6 μl of DNA library after the first 6 hours of the sequencing run. Base-calling was performed using the Metrichor cloud software and all ONT reads (quality passed 1D and 2D) as well as only 2D reads were then extracted using Poretools^43^ for downstream analysis.

### Illumina library preparation and sequencing

The sequencing libraries for all clinical isolates (n=14) were prepared using Nextera XT DNA library preparation kit (Illumina, San Dego, CA, USA) and sequenced on the MiSeq Platform (Illumina), according to manufacturer’s instructions, resulting in an average of 967,369 reads with an average read length of 257 bp after quality control and average coverage of 86.5 × per base. The raw sequence files for the 2016 WHO gonococcal reference strains^41^ were obtained from the European Nucleotide Archive (ENA; Bioproject PRJEB14020). The 2D ONT reads and Illumina reads were species confirmed using the online tool One Codex (www.onecodex.com). All sequenced reads are available from the European Nucleotide Archive with the following accession number: PRJEB25703.

### Assembly and assessment

To identify the ideal tool for obtaining a high quality *de novo* assembly using only ONT reads, we evaluated Canu (v1.6)^44^, Miniasm (vr122)^45^, PBcR (v8.3)^46^, and SMARTdenovo (available from https://github.com/ruanjue/smartdenovo) for assembly of all ONT reads (1D and 2D) and only the extracted 2D ONT reads. Canu was executed with default parameters including error correction and --genomesize 2.2M. Miniasm was run with default parameters. PBcR was executed with the following parameters: -length 500, -partitions 200 and –genomeSize=2200000. SMARTdenovo was run with default parameters and –c 1 to run consensus step. Subsequently, hybridSPAdes (v3.11.1)^47^ and MaSuRCA (v3.2.2)^48^ were used separately to produce hybrid assemblies by including paired-end Illumina reads in the assembly process. hybridSPAdes was executed with the careful option and –nanopore command. MaSuRCA was performed using the following parameters: GRAPH_KMER_SIZE=auto, USE_LINKING_MATES=0, LIMIT_JUMP_COVERAGE=60, CA_PARAMETERS=cgwErrorRate=0.25, NUM_THREADS=64, JF_SIZE=23000000, DO_HOMOPOLYMER_TRIM=0.

The assemblies obtained from Miniasm using 2D ONT reads were error corrected twice using Racon (v1.3.0)^49^ and polished with the signal-level consensus software, nanopolish (v0.7.1) (available from: https://github.com/jts/nanopolish). Briefly, 2D ONT reads were mapped as all-vs-all read self-mapping using Minimap with the following options: -S, -w 5, -L 100, -m 0. Subsequently, Miniasm was executed with default options. The 2D ONT reads were mapped back to the assembly with Minimap (default parameters) and corrected with Racon with default parameters, and this was repeated one time. Burrows-Wheeler Aligner (v0.7.17-r1188) (BWA-MEM)^50^ with the ’x ont2d option was then used to map all reads back to the Racon-corrected assembly to be used as the input for Nanopolish. Nanopolish was run with default parameters and --min-candidate-frequency 0.1. Furthermore, Canu assemblies were error corrected with Pilon (v1.22)^51^, Circlator (v1.5.3)^52^ and Nanopolish. All error corrections for the Canu assemblies with the above-mentioned tools were performed with default parameters.

Finally, to evaluate the accuracy of our assemblies, the ONT (corrected and noncorrected) and hybrid assemblies for the WHO reference strains were compared with the finished and closed 2016 WHO gonococcal reference genomes^41^. All 2D ONT reads were aligned to the genome sequence of respective WHO reference strain, to be able to easily and directly determine the quality and accuracy using BWA-MEM (for mapping and phylogenetics) and Quast (v4.6.0) (for *de novo* assemblies). The top three assessed ONT *de novo* assemblies and the best hybrid assemblies were subjects for downstream analysis. Gene coding sequences (CDS) were annotated using Prokka (v1.12) on the chosen assemblies^53^.

### Multiple-sequence alignment and phylogenetics

All Illumina and 2D ONT reads were mapped to the chromosome of FA1090 (GenBank: NC_002946.2) separately using BWA-MEM, the 2D ONT reads were specified using the nanopore option (-x –ont2d). All consensus sequences were merged into a multiple sequence alignment, single-nucleotide polymorphisms (SNPs) were called, and a maximum-likelihood phylogenetic tree based on SNPs was obtained using RAxML (version 8.2.8)^54^. This was also performed on Illumina sequences and 2D ONT sequences separately and the phylogeny was compared in a tanglegram.

### Antimicrobial resistance determinants

To detect AMR determinants relevant for gonococci as quick and straight-forwardly as possible for future routine use of MinION sequencing, we evaluated several open-access and user-friendly web-based sequence analysis tools. The top three assessed ONT assemblies and the best hybrid assembly were examined using the Whole Genome Sequence Analysis (WGSA; www.wgsa.net)^21^ and the PubMLST *N. gonorrhoeae* AMR (still in development) scheme (www.pubmlst.org)^55^. The focus was on AMR determinants for ceftriaxone and cefixime (*penA*, *mtrR*, *penB*), azithromycin (*23S rDNA*, *mtrR*), ciprofloxacin (gyrA), tetracycline (*rpsJ*, *tet(M)* (plasmid-mediated resistance)), and penicillin (*penA*, *ponA*, *mtrR*, *penB*, *bla*_TEM_ (plasmid-mediated resistance)). We only detected the AMR determinants on the plasmids and no additional downstream analysis was performed on the plasmids.

Furthermore, the 2D ONT reads were examined in our *in house*-customised CLC Genomics Workbench workflow to characterise the AMR determinants based on alleles in the *N. gonorrhoeae* Sequence Typing for Antimicrobial Resistance (NG-STAR) database (www.ngstar.canada.ca)^56^. The NG-STAR database lacked the possibility to run contig assemblies at the time of this study. The workflow employed *de novo* assembly without mapping and used BLAST for the identification, with the highest (%) hit being reported^57^.

The best performing tool of assembly and subsequent detection of AMR determinants was subsequently used to characterise the clinical isolates (n=14).

## Results

### Overview of sequenced data

The 14 2016 WHO reference strains^41^ and 14 clinical isolates were sequenced using the ONT MinION platform. In these MinION runs, we obtained between 105 Mb and 922 Mb of 2D sequences with longest reads ranging from 26.0 to 58.6 kb and an average read length of 4.6-6.4 kb. The number of passed 2D ONT reads ranged from 17,500 to 147,700 with an average of 33% of all reads. For evaluation of the ONT sequences and further downstream analysis (taxonomy, hybrid assemblies and phylogeny), we used the 2×100 bp Illumina HiSeq paired-end reads with >250 depth for each WHO reference genome^41^ and 2×300 bp Illumina MiSeq paired-end reads with 80-100× depth for each clinical isolate. Taxonomy was defined for all reads using One Codex and 97-99% and 100% of all reads were classified as *N. gonorrhoeae* using ONT (Fig. 1) and Illumina MiSeq, respectively.

**Figure 1.**
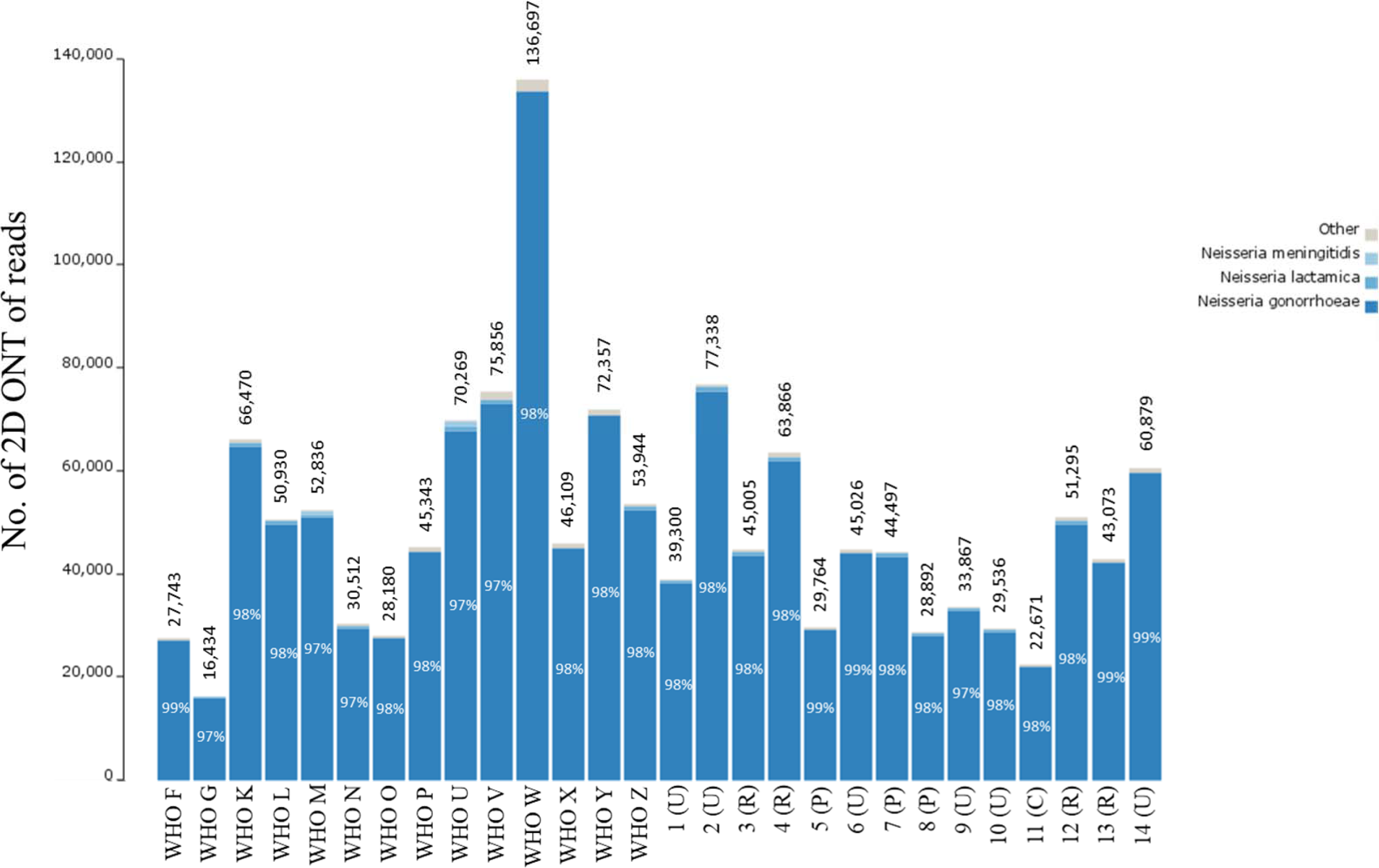
Readcount for the MinION (Oxford Nanopore Technologies (ONT)) dataset and read taxonomy classification of 2D reads, when sequencing the 2016 WHO *Neisseria gonorrhoeae* reference strains (n=14)^41^ and 14 clinical *N. gonorrhoeae* isolates. Percentage within each bar represents the proportion of reads classified as *N. gonorrhoeae.* The number above each bar represents the number of reads that were classified.

### Comparison of MinION *de novo* assemblies with the 2016 WHO gonococcal reference genomes^41^

Assemblies were compared and selected based on the Quast results when comparing to the WHO reference genomes using the following criteria: lowest number of mismatches, misassemblies and number of contigs, and highest fraction of genome covered (Supplementary Table). We selected and analysed ONT assemblies produced by Canu using only 2D reads and polished with Nanopolish, Miniasm twice error corrected with Racon and polished with Nanopolish, and SMARTdenovo. Furthermore, the MaSuRCA hybrid assemblies (based on the ONT plus Illumina reads) were selected for downstream analysis (Table 1). Generally, ONT assemblies based on only the 2D ONT reads had a higher accuracy compared with assemblies using all reads (1D and 2D ONT reads) with fewer mismatches but very modest to no change in contiguity of the assemblies and fraction of the WHO reference genome covered. All error corrections were made using only 2D ONT reads and statistics such as mismatches and indels improved. Nanopolish had the best performance of the tested error correction tools and reduced the number of indels by up to 13 times. Assemblies produced with hybridSPAdes were more affected by the lower numbers of ONT reads, i.e. when using only 2D ONT reads. The hybrid assembly using SPAdes had less contiguity (up to 5.5 times more contigs) with higher number of mismatches (up to 4.4 more mismatches per 100 kb) and introductions of misassemblies (Supplementary Table). In general, the 2D ONT reads were used more efficiently by MaSuRCA for hybrid assembly, with overall improved statistics especially for contiguity and mismatches.

**Table 1.** Statistics for the three chosen MinION (Oxford Nanopore Technologies)-only *de novo* assemblies and the MinION-Illumina hybrid assembly, when sequencing the 2016 WHO *Neisseria gonorrhoeae* reference strains (n=14)^41^.

|  |  | WHO F | WHO G | WHO K | WHO L | WHO M | WHO N | WHO O | WHO P | WHO U | WHO V | WHO W | WHO X | WHO Y | WHO Z |
| --- | --- | --- | --- | --- | --- | --- | --- | --- | --- | --- | --- | --- | --- | --- | --- |
| Canu<sup>44</sup> +<br>Nanopolish | Genome fraction (%) | 98.96 | 99.99 | 100.00 | 99.97 | 99.59 | 100.00 | 99.99 | 99.96 | 99.81 | 100.00 | 89.93 | 100.00 | 100.00 | 99.48 |
|  | Contigs | 1 | 5 | 3 | 3 | 2 | 3 | 3 | 1 | 1 | 1 | 1 | 1 | 1 | 1 |
|  | Largest contig (bp) | 2266147 | 1611568 | 1415719 | 2166845 | 2178610 | 2182677 | 2178602 | 2170535 | 2244376 | 2231681 | 1997317 | 2190700 | 2234645 | 2222403 |
|  | Mismatches / 100 kb | 301.10 | 205.07 | 194.76 | 278.69 | 226.92 | 146.03 | 258.56 | 266.96 | 186.77 | 113.13 | 277.04 | 190.09 | 268.69 | 196.64 |
|  | Indels / 100 kb | 309.39 | 412.92 | 182.73 | 261.07 | 207.51 | 208.12 | 255.61 | 257.39 | 158.88 | 92.15 | 239.01 | 191.75 | 320.37 | 265.95 |
|  | Annotated CDS | 3893 | 4172 | 3632 | 3783 | 3717 | 3898 | 3770 | 3561 | 3726 | 3639 | 3210 | 3543 | 3863 | 3850 |
| Miniasm<sup>45</sup> +<br>Racon<sup>49</sup> +<br>Nanopolish | Genome fraction (%) | 99.48 | 99.96 | 100.00 | 100.00 | 100.00 | 95.67 | 99.99 | 99.89 | 100.00 | 99.99 | 99.97 | 100.00 | 99.99 | 98.22 |
|  | Contigs | 7 | 6 | 6 | 9 | 7 | 6 | 4 | 7 | 6 | 3 | 7 | 1 | 1 | 71 |
|  | Largest contig (bp) | 761198 | 744021 | 1060644 | 1288971 | 938255 | 2063596 | 2168124 | 1621005 | 1775729 | 2220894 | 1171322 | 2171999 | 2229595 | 1847314 |
|  | Mismatches / 100 kb | 249.03 | 262.03 | 161.07 | 291.29 | 107.47 | 310.14 | 290.36 | 198.44 | 229.38 | 265.48 | 119.6 | 256 | 340.97 | 237.81 |
|  | Indels / 100 kb | 275.87 | 616.64 | 394.5 | 448.49 | 120.41 | 497.18 | 393.58 | 495.94 | 276.87 | 327.2 | 140.89 | 255.68 | 358.28 | 238.45 |
|  | Annotated CDS | 3997 | 4485 | 4321 | 4276 | 3789 | 4240 | 3999 | 4418 | 4146 | 3848 | 3967 | 3515 | 3969 | 4373 |
| SMARTdenovo | Genome fraction (%) | 99.02 | 100.00 | 100.00 | 100.00 | 100.00 | 99.98 | 100.00 | 100.00 | 99.62 | 100.00 | 100.00 | 100.00 | 100.00 | 100.00 |
|  | Contigs | 2 | 2 | 2 | 2 | 3 | 3 | 3 | 1 | 15 | 1 | 2 | 1 | 1 | 1 |
|  | Largest contig (bp) | 1749604 | 2152447 | 1769445 | 2167238 | 2182509 | 2162509 | 2160811 | 2167730 | 1026600 | 2213205 | 2213608 | 2163199 | 2210497 | 2211396 |
|  | Mismatches / 100 kb | 219.03 | 111.24 | 97.38 | 124.36 | 179.05 | 202.09 | 173.35 | 114.91 | 224.47 | 26.65 | 226.22 | 175.3 | 246.44 | 237.6 |
|  | Indels / 100 kb | 680.4 | 822.11 | 670.05 | 690.99 | 654.08 | 686.08 | 662.68 | 669.96 | 657.5 | 641.03 | 659.5 | 639.12 | 817.77 | 735.46 |
|  | Annotated CDS | 4602 | 4470 | 4388 | 4523 | 4512 | 4614 | 4526 | 4372 | 4712 | 4378 | 4301 | 4342 | 4553 | 4461 |
| MaSuRCA<sup>48</sup> | Genome fraction (%) | 99.46 | 99.99 | 99.91 | 99.99 | 99.97 | 99.99 | 99.99 | 99.99 | 100.00 | 99.99 | 99.99 | 99.99 | 99.99 | 99.95 |
|  | Contigs | 1 | 3 | 3 | 2 | 5 | 3 | 5 | 3 | 2 | 7 | 8 | 6 | 4 | 1 |
|  | Largest contig (bp) | 2279985 | 2176816 | 2128117 | 2190405 | 2189696 | 2188414 | 2197012 | 2204260 | 2258223 | 2236154 | 2238072 | 2171685 | 2248815 | 2255505 |
|  | Mismatches / 100 kb | 1.89 | 0.74 | 2.54 | 3.97 | 1.38 | 2.12 | 2.31 | 2.48 | 2.55 | 0.14 | 0.95 | 2.9 | 0.45 | 3.68 |
|  | Indels / 100 kb | 2.06 | 1.02 | 1.71 | 1.38 | 1.01 | 1.66 | 1.71 | 1.89 | 1.61 | 1.35 | 2.43 | 1.98 | 0.99 | 1.88 |
|  | Annotated CDS | 2304 | 2229 | 2228 | 2261 | 2307 | 2264 | 2306 | 2241 | 2312 | 2362 | 2400 | 2230 | 2333 | 2259 |

CDS, coding sequence

Although the length of the assemblies based on the ONT sequences did not largely differ from the WHO reference genomes, the number of CDS was vastly different. The 2016 WHO gonococcal reference genomes include 2295-2450 CDS^41^. However, 3210-4712 CDS were identified using Prokka in our selected ONT assemblies. In contrast, Prokka identified 2228-2400 CDS in the MaSuRCA hybrid assemblies, which is in line with the published WHO reference genomes (Table 1).

Error corrections using Pilon and Circlator, which is not designed for error correction, did not substantially improve the assemblies (Supplementary Table) and the assemblies error corrected with these tools were not used in the downstream analysis.

### MinION for molecular epidemiology, including rapid outbreak investigations

The Illumina reads mapped to the PacBio-produced WHO reference genomes (n=14)^41^ with no SNPs detected except for WHO M (one SNP) and a range of 93.3% to 96.4% of the reads mapped to the respective PacBio reference genome. The ONT reads mapped with 2-58 SNPs per genome sequence and 95.2% to 98.3% of the reads mapped. By executing BWA-MEM with the nanopore -x ont2d option, no SNPs between the ONT reads and the respective reference genomes were detected, but we found variation in the number of 2D ONT reads and percentage mapped to the FA1090 reference genome (Fig. 2). Therefore, the BWA-MEM nanopore option was used when mapping all 2D ONT reads to the FA1090 reference genome for the phylogenetic analysis (Fig. 3). By creating the phylogeny with the ONT-produced genomes (n=28) and Illumina MiSeq genomes (n=28) separately, we showed that the ONT sequenced libraries produced a phylogenetic tree topology that was comparable with the one using the Illumina dataset (Fig. 3b). The main difference in the tree topology was that WHO G and WHO N were closer to the root when Illumina sequences were used for the phylogeny. Moreover, all identical isolates separately sequenced with ONT and Illumina clustered together, except for some (n=7) of the clinical isolates where the Illumina sequences were more closely related. In four of these cases (11 cervix (C)/12 rectum (R) and 13 (R)/14 urethra (U)), the isolates were from the same patients but from different anatomical sites. For three isolates (5, 6 and 7), the Illumina sequences were also more closely related but all of the separately sequenced datasets were still on the same branch showing that these isolates were very similar (Fig. 3a).

**Figure 2.**
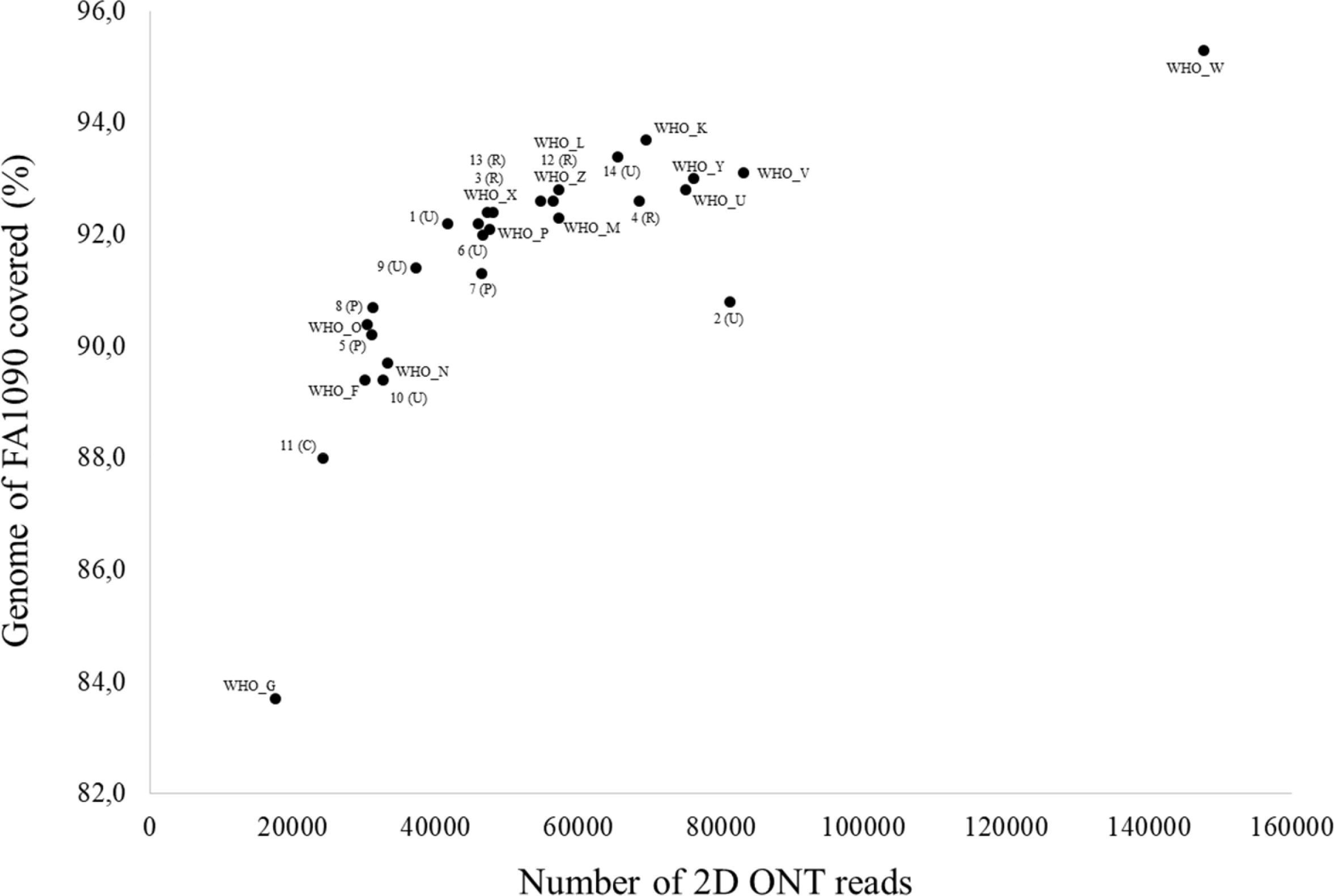
Overview of the number of MinION (Oxford Nanopore Technologies (ONT)) 2D reads and the mapability of the reads to the genome of the *Neisseria gonorrhoeae* reference strain FA1090, when sequencing the 2016 WHO *N. gonorrhoeae* reference strains (n=14)^41^ and 14 clinical *N. gonorrhoeae* isolates.

**Figure 3.**
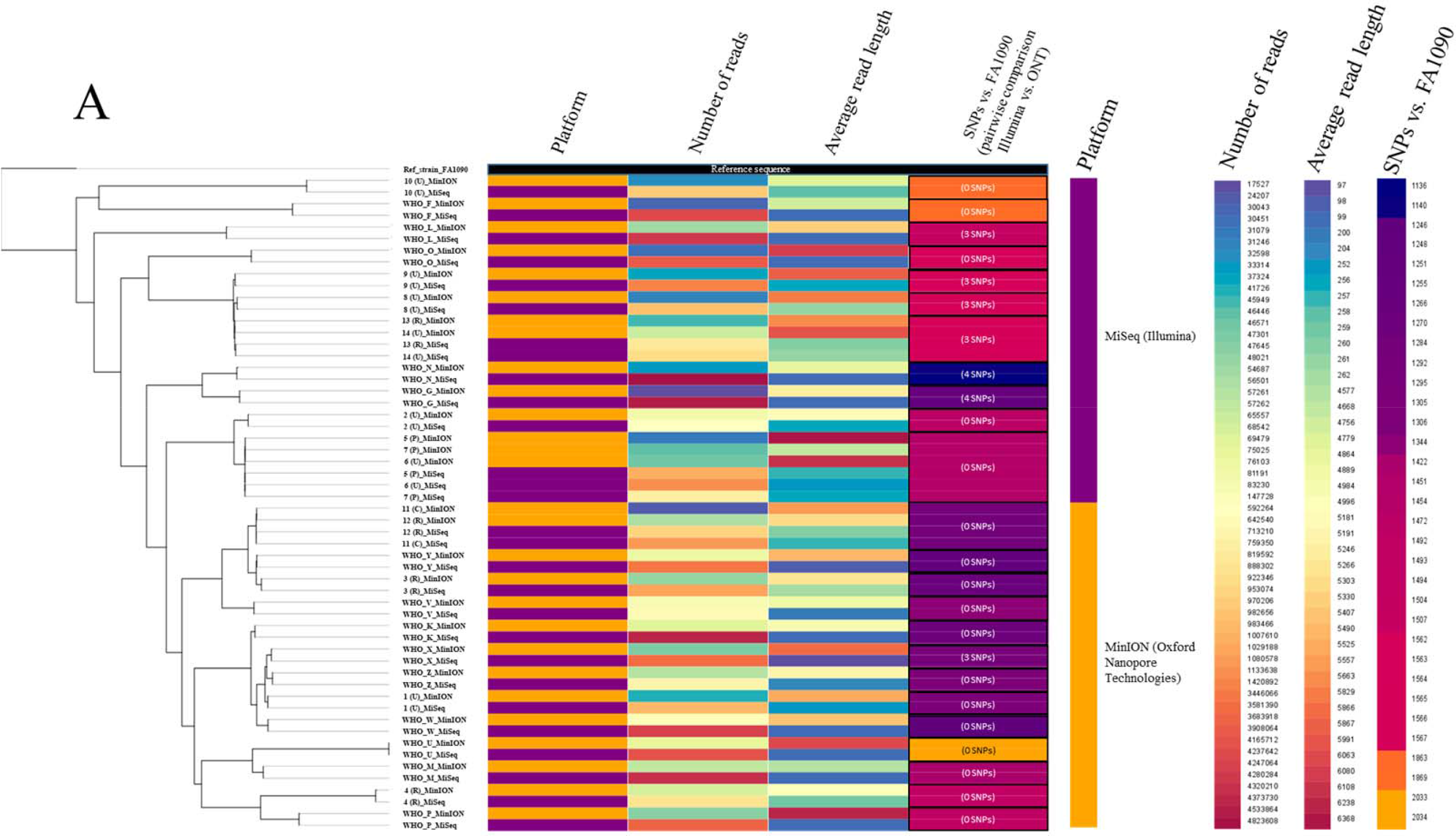
(A) Phylogenetic tree of the genome sequences of the 2016 WHO *N. gonorrhoeae* reference strains (n=14)^41^ and clinical gonococcal isolates (n=14) sequenced with Illumina MiSeq and MinION (Oxford Nanopore Technologies (ONT)). The tree uses using the genome of the *N. gonorrhoeae* reference strain FA1090 as reference (shown with black bar). The platform used, number of reads, average read length, and number of single nucleotide polymorphisms (SNPs) are displayed as colored bars next to each node in the tree. The numbers inside the SNP-bars is the pairwise distance between the Illumina and ONT sequences. (B) Tanglegram to compare the phylogenetic networks based on sequencing using Illumina technology (left hand side) and Oxford Nanopore technology (right hand side).

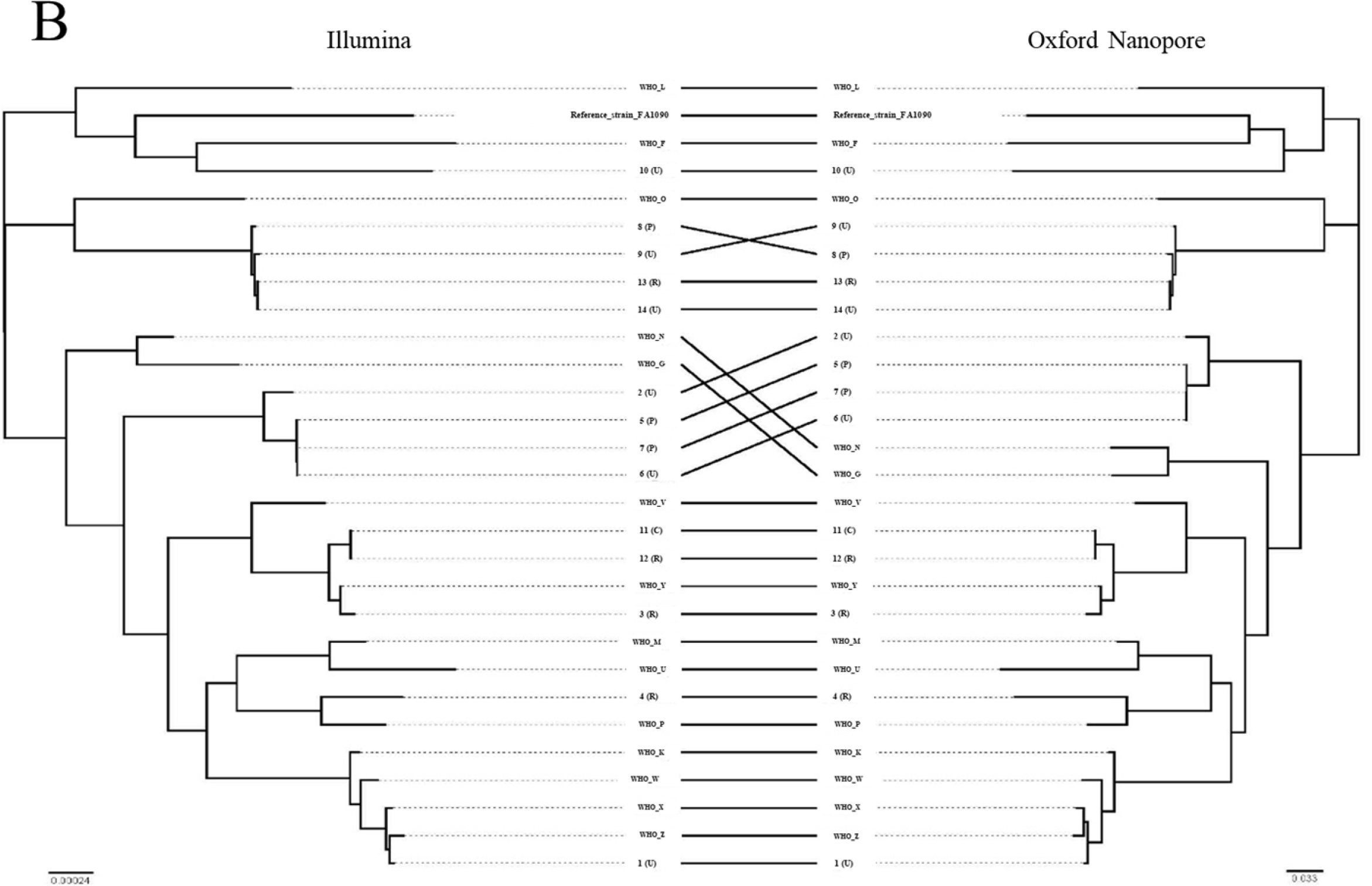

### MinlON for detection of AMR determinants

To investigate the four final *de novo* assemblies using Canu (polished), Miniasm (polished), SMARTdenovo, and MaSuRCA (Illumina hybrid) for detection of AMR determinants in the 2016 WHO gonococcal reference genomes^41^, we used the WGSA (www.wgsa.net)^21^ and PubMLST (www.pubmlst.org).

WGSA is an easy-to-use web interface where you drag-and-drop the assemblies into the web browser and the characterisation is done in minutes^21^. We used this interface for all chosen assemblies (n=112) and the analysis of AMR determinants took 15 minutes. We observed good concordance overall with the verified AMR determinants in the reference genomes (Table 2), with the exception of the 23S *rDNA* macrolide resistance determinant C2611T, which was incorrectly identified in the Canu (polished) and Miniasm (polished) assemblies. No ONT assembly was able to correctly characterise *penA* in WHO W and none, including the MaSuRCA hybrid assembly, was able to correctly characterise *penA* in WHO K (Table 2). The characterisation of plasmid-mediated AMR determinants was inaccurate in all four chosen assemblies for the detection of *tet(M)*. Also, the chosen assemblies failed to detect *bla*_TEM_ in WHO M, WHO O and WHO V and incorrectly reported presence of *bla*_TEM_ in WHO G in the Miniasm (polished), SMARTdenovo and MaSuRCA assemblies (Table 3). Out of the 2D ONT assemblies, the SMARTdenovo assembly performed the best.

**Table 2.** Characterisation of chromosomal antimicrobial resistance (AMR) determinants in the 2016 WHO *Neisseria gonorrhoeae* reference strains (n=14)^41^ using Whole Genome Sequence Analysis (WGSA; www.wgsa.net)^21^ and three MinION-only assemblies, MinION-Illumina hybrid assembly and an *in house*-customised CLC Genomics Workbench, including NG-STAR56 and a customised database for AMR determinant detection. The discrepant results from the published reference genomes are in bold. The correct AMR determinants are in italic.

|  |  |  | WHO F | WHO G | WHO K | WHO L | WHO M | WHO N | WHO O | WHO P | WHO U | WHO V | WHO W | WHO X | WHO Y | WHO Z |
| --- | --- | --- | --- | --- | --- | --- | --- | --- | --- | --- | --- | --- | --- | --- | --- | --- |
| wgsa.net | Canu + Nanopolish<sup>44</sup> | penA | WT | WT | WT | A501V | WT | WT | WT | WT | WT | WT | WT | Mosaic | Mosaic, A501P | Mosaic |
|  |  | mtrR (A-deletion/G45D) | No | No | Yes | Yes | Yes | No | No | No | No | Yes | No | Yes | Yes | No |
|  |  | penB (G101, A102) | WT, WT | WT, WT | K, D | K, D | K, D | WT, WT | K, D | WT, D | WT, WT | K, D | K, D | K, D | K, N | K, D |
|  |  | 23S rDNA | C2611T | WT | WT | WT | WT | WT | WT | C2611T | C2611T | A2059G | WT | C2611T | C2611T | C2611T |
|  |  | gyrA (S91F) | No | Yes | Yes | Yes | Yes | Yes | No | No | No | Yes | Yes | Yes | Yes | Yes |
|  |  | ponA (L421P) | No | Yes | Yes | Yes | Yes | Yes | Yes | No | Yes | Yes | Yes | Yes | Yes | Yes |
|  |  | rpsJ (V57M) | No | Yes | Yes | Yes | Yes | Yes | Yes | Yes | Yes | Yes | Yes | Yes | Yes | Yes |
|  | Miniasm + Racon + Nanopolish<sup>45,49</sup> | penA | WT | WT | WT | A501V | WT | WT | WT | WT | WT | WT | WT | Mosaic | A501P | Mosaic |
|  |  | mtrR (A-deletion/G45D) | No | No | Yes | Yes | Yes | No | No | No | No | Yes | Yes | Yes | Yes | No |
|  |  | penB (G101,A102) | WT, WT | WT, WT | K, D | K, D | K, D | WT, WT | K, D | WT, D | WT, WT | K, D | K, D | K, D | K, N | K, D |
|  |  | 23S rDNA | C2611T | WT | WT | WT | C2611T | WT | WT | WT | C2611T | A2059G | WT | WT | WT | WT |
|  |  | gyrA (S91F) | No | Yes | Yes | Yes | Yes | Yes | No | No | No | Yes | Yes | Yes | Yes | Yes |
|  |  | ponA (L421P) | No | Yes | Yes | Yes | Yes | Yes | Yes | No | Yes | Yes | Yes | Yes | Yes | Yes |
|  |  | rpsJ (V57M) | No | Yes | Yes | Yes | Yes | Yes | Yes | Yes | Yes | Yes | Yes | Yes | Yes | Yes |
|  | SMARTdenovo | penA | WT | WT | WT | A501V | WT | WT | WT | WT | WT | WT | WT | Mosaic | A501P | Mosaic |
|  |  | mtrR (A-deletion/G45D) | No | No | Yes | Yes | Yes | No | No | No | No | No | No | No | No | No |
|  |  | penB (G101,A102) | WT, WT | WT, WT | K, D | K, D | K, D | WT, WT | K, D | WT, D | WT, WT | K, D | K, D | K, D | K, N | K, D |
|  |  | 23S rDNA | WT | WT | WT | WT | WT | WT | WT | WT | C2611T | A2059G | WT | WT | WT | WT |
|  |  | gyrA (S91F) | No | Yes | Yes | Yes | Yes | Yes | No | No | No | Yes | Yes | Yes | Yes | Yes |
|  |  | ponA (L421P) | No | Yes | Yes | Yes | Yes | Yes | Yes | No | Yes | Yes | Yes | Yes | Yes | Yes |
|  |  | rpsJ (V57M) | No | No | Yes | Yes | Yes | Yes | Yes | Yes | Yes | Yes | Yes | Yes | Yes | Yes |
|  | MaSuRCA<sup>48</sup> | penA | WT | WT | WT | A501V | WT | WT | WT | WT | WT | WT | WT | Mosaic | A501P | Mosaic |
|  |  | mtrR (A-deletion/G45D) | No | Yes | Yes | Yes | Yes | No | Yes | No | No | Yes | Yes | Yes | Yes | No |
|  |  | penB (G101,A102) | WT, WT | WT, WT | K, D | K, D | K, D | WT, WT | K, D | WT, D | WT, WT | K, D | K, D | K, D | K, N | K, D |
|  |  | 23S rDNA | WT | WT | WT | WT | WT | WT | WT | WT | C2611T | A2059G | WT | WT | WT | WT |
|  |  | gyrA (S91F) | No | Yes | Yes | Yes | Yes | Yes | No | No | No | Yes | Yes | Yes | Yes | Yes |
|  |  | ponA (L421P) | No | Yes | Yes | Yes | Yes | Yes | Yes | No | Yes | Yes | Yes | Yes | Yes | Yes |
|  |  | rpsJ (V57M) | No | Yes | Yes | Yes | Yes | Yes | Yes | Yes | Yes | Yes | Yes | Yes | Yes | Yes |
|  | In house-customised CLC Genomics Workbench | penA | WT | WT | Mosaic | A501V | WT | WT | WT | WT | WT | WT | Mosaic | Mosaic | Mosaic, A501P | Mosaic |
|  |  | mtrR (A-deletion/G45D) | No | Yes | Yes | Yes | Yes | No | Yes | No | No | Yes | Yes | Yes | Yes | No |
|  |  | penB (G101,A102) | WT, WT | WT, WT | K, D | K, D | K, D | WT, WT | K, D | WT, D | WT, WT | K, D | K, D | K, D | K, N | K, D |
|  |  | 23S rDNA | WT | WT | WT | WT | WT | WT | WT | WT | C2611T | A2059G | WT | WT | WT | WT |
|  |  | gyrA (S91F) | No | Yes | Yes | Yes | Yes | Yes | No | No | No | Yes | Yes | Yes | Yes | Yes |
|  |  | ponA (L421P) | No | Yes | Yes | Yes | Yes | Yes | Yes | No | Yes | Yes | Yes | Yes | Yes | Yes |
|  |  | rpsJ (V57M) | No | Yes | Yes | Yes | Yes | Yes | Yes | Yes | Yes | Yes | Yes | Yes | Yes | Yes |

**Table 3.** Detection of plasmid-mediated antimicrobial resistance determinants in the 2016 WHO *Neisseria gonorrhoeae* reference strains (n=14)^41^ using Whole Genome Sequence Analysis (WGSA; www.wgsa.net)^21^ and an *in house*-customised CLC Genomics Workbench. The discrepant results from the published references are in bold.

|  |  |  | WHO F | WHO G | WHO K | WHO L | WHO M | WHO N | WHO O | WHO P | WHO U | WHO V | WHO W | WHO X | WHO Y | WHO Z |
| --- | --- | --- | --- | --- | --- | --- | --- | --- | --- | --- | --- | --- | --- | --- | --- | --- |
| www.wgsa.net | <sup>44</sup> Canu + Nanopolish | bla<sub>TEM</sub> | No | No | No | No | **No** | Yes | **No** | No | No | **No** | No | No | No | No |
|  |  | tet(M) | No | **No** | No | No | No | **No** | No | No | No | No | No | No | No | No |
|  | <sup>45</sup> Miniasm +<br><sup>49</sup> Racon + Nanopolish | bla<sub>TEM</sub> | No | **Yes** | No | No | **No** | Yes | **No** | No | No | **No** | No | No | No | No |
|  |  | tet(M) | No | **No** | No | No | No | **No** | No | No | No | No | No | No | No | No |
|  | SMARTdenovo | bla<sub>TEM</sub> | No | **Yes** | No | No | **No** | Yes | **No** | No | No | **No** | No | No | No | No |
|  |  | tet(M) | No | **No** | No | No | No | **No** | No | No | No | No | No | No | No | No |
|  | <sup>48</sup> MaSuRCA | bla<sub>TEM</sub> | No | **Yes** | No | No | **No** | Yes | **No** | No | No | **No** | No | No | No | No |
|  |  | tet(M) | No | **No** | No | No | No | **No** | No | No | No | No | No | No | No | No |
|  | In house-customised<br>CLC<br>Genomics<br>Workbench | bla<sub>TEM</sub> | No | No | No | No | Yes | Yes | Yes | No | No | Yes | No | No | No | No |
|  |  | tet(M) | No | Yes | No | No | No | Yes | No | No | No | No | No | No | No | No |
|  | Publis<br>hed<br>referen<br>ce<sup>41</sup> | bla<sub>TEM</sub> | No | No | No | No | Yes | Yes | Yes | No | No | Yes | No | No | No | No |
|  |  | tet(M) | No | Yes | No | No | No | Yes | No | No | No | No | No | No | No | No |

WT, Wild type.

We also used the PubMLST database, which is another easy-to-use web based tool, that can mine assembly data for various purposes. The database uses assemblies as input and delivers results in few minutes. The PubMLST database (www.pubmlst.org), using 2D ONT assemblies, was only able to characterise the *mtrR* AMR determinant and this was only done in the Canu assemblies. However, using the MaSuRCA hybrid assembly PubMLST had 100% concordance with the published reference genomes for detection of *penA*, *mtrR*, *penB*, *ponA*, *23S rDNA*, *gyrA*, and *tet(M)*. PubMLST does not include *rpsJ* in its gonococcal AMR characterisation module and also failed to detect the *bla*_TEM_ gene in WHO M and WHO N (Table 3).

The NG-STAR database^56^, included in the WHO CC *in house*-customised CLC Genomics Workbench, gave 100% concordance for the presence of all AMR determinants characterised in NG-STAR^56^ with the reference genomes (Table 2). All the plasmid-mediated AMR determinants (Table 3) and the AMR determinants *penB*, *16S rDNA*, and *parC* were also correctly characterised. The customised CLC Genomics Workbench was therefore used to characterise the AMR determinants in the clinical isolates using only the 2D ONT reads.

### *In* AoMse-customised CLC Genomics Workbench for characterisation of AMR determinants in clinical gonococcal isolates

The 14 isolates expressed decreased susceptibility (MIC≥0.032 mg/L)^58^ or resistance to ceftriaxone (11/14) and cefixime (11/14), azithromycin (3/14), and ciprofloxacin (13/14) (Table 4). No isolate displayed spectinomycin resistance or produced β-lactamase. The *in house*-customised CLC Genomics Workbench including the NG-STAR database^56^ correctly characterised all AMR determinants in the isolates using only the 2D ONT reads and found 12, 1, and 13 of the 14 isolates carrying AMR mutations in *penA* (cefixime and ceftriaxone resistance determinant), *23S rDNA* (azithromycin resistance determinant), and *gyrA* (ciprofloxacin resistance determinant), respectively. Twelve of the 14 isolates also harboured *parC* resistance mutations, which contribute to the high MICs in isolates with high-level ciprofloxacin resistance. Specific mutations in *mtrR*, resulting in an over-expression of the MtrCDE efflux pump, and *porB1b* (*penB* AMR determinant), causing a decreased influx of antimicrobials through PorB, were identified in 12 of the 14 isolates. No *16S rDNA* resistance mutations (spectinomycin resistance determinant) or plasmid-mediated AMR determinants (high-level resistance to penicillin and tetracycline) were detected. Resistance mutation in *rpsJ* (decreased susceptibility/resistance to tetracycline) was found in all isolates. There was 100% concordance between the AMR determinants detected using the 2D ONT reads only and Illumina reads. Consequently, resistance or decreased susceptibility to extended-spectrum cephalosporins, azithromycin, spectinomycin, ciprofloxacin, tetracycline, and penicillin could be accurately predicted using solely 2D ONT reads (Table 4).

**Table 4.** Minimum inhibitory concentrations (MIC, mg/L) of seven antimicrobials in 14 clinical *Neisseria gonorrhoeae* isolates and associated antimicrobial resistance determinants characterised using an *in house*-customised CLC Genomics Workbench with MinION (Oxford Nanopore Technology) reads.

|  |  | 1<br>(U) | 2<br>(U) | 3<br>(R) | 4<br>(R) | 5<br>(P) | 6<br>(U) | 7<br>(P) | 8<br>(P) | 9<br>(U) | 10<br>(U) | 11<br>(C) | 12<br>(R) | 13<br>(R) | 14<br>(U) |
| --- | --- | --- | --- | --- | --- | --- | --- | --- | --- | --- | --- | --- | --- | --- | --- |
| MIC interpretation<br>(S/I/R)<sup>a</sup> | Ceftriaxone (MIC, mg/L) | S (0.032) | S (0.064) | S (0.016) | S (0.002) | S (0.064) | S (0.064) | S (0.064) | S (0.064) | S (0.032) | S (0.004) | S (0.032) | S (0.032) | S (0.064) | S (0.064) |
|  | Cefixime (MIC, mg/L) | R (0.25) | S (0.064) | S (0.064) | S (<0.016) | S (0.064) | S (0.032) | S (0.032) | S (0.064) | S (0.016) | S (<0.016) | S (0.125) | S (0.125) | S (0.064) | S (0.064) |
|  | Azithromycin (MIC, mg/L) | S (0.125) | S (0.25) | I (0.5) | I (0.5) | S (0.25) | S (0.5) | S (0.25) | S (0.25) | S (0.5) | R (4) | R (1) | R (1) | I (0.5) | I (0.5) |
|  | Ciprofloxacin (MIC, mg/L) | R (>32) | R (4) | R (>32) | R (0.125) | R (16) | R (32) | R (16) | R (>32) | R (>32) | S (0.004) | R (>32) | R (>32) | R (>32) | R (>32) |
|  | Spectinomycin (MIC, mg/L) | S (16) | S (8) | S (8) | S (8) | S (16) | S (16) | S (16) | S (8) | S (16) | S (16) | S (16) | S (16) | S (16) | S (16) |
|  | Penicillin (MIC, mg/L) | I (1) | R (2) | R (2) | I (1) | R (4) | R (4) | R (2) | R (2) | R (2) | I (0.5) | R (2) | R (2) | R (2) | R (2) |
|  | Tetracycline (MIC, mg/L) | I (1) | I (1) | R (2) | S (0.25) | R (2) | R (2) | I (1) | S (0.5) | S (0.25) | S (0.25) | I (1) | I (1) | S (0.5) | S (0.5) |
| Antimicrobial resistance determinants<br>extracted using an *in house*-customised CLC<br>Genomics Workbench | *penA* (Mosaic/A501V/T) | Mosaic | A501V | Mosaic | WT | A501V | A501V | A501V | A501T | A501T | WT | Mosaic | Mosaic | A501T | A501T |
|  | *mtrR* (A-deletion/G45D) | Yes | Yes | Yes | No | Yes | Yes | Yes | Yes | Yes | No | Yes | Yes | Yes | Yes |
|  | *penB* variant | Yes | Yes | Yes | WT | Yes | Yes | Yes | Yes | Yes | WT | Yes | Yes | Yes | Yes |
|  | 23S *rDNA* (A2059, C2611) | WT | WT | WT | WT | WT | WT | WT | WT | WT | C2611T | WT | WT | WT | WT |
|  | 16S *rDNA* (C1194T) | No | No | No | No | No | No | No | No | No | No | No | No | No | No |
|  | *gyrA* (S91F) | Yes | Yes | Yes | Yes | Yes | Yes | Yes | Yes | Yes | No | Yes | Yes | Yes | Yes |
|  | *parC* (S86, S87, E91) | S87R | D86N | S87R | WT | D86N | D86N | D86N | E91G | E91G | WT | S87R | S87R | E91G | E91G |
|  | *ponA* (L421P) | Yes | Yes | Yes | No | Yes | Yes | Yes | Yes | Yes | No | Yes | Yes | Yes | Yes |
|  | *penA* (D345a) | No | Yes | No | Yes | Yes | Yes | Yes | Yes | Yes | Yes | No | No | Yes | Yes |
|  | *rpsJ* (V57M) | Yes | Yes | Yes | Yes | Yes | Yes | Yes | Yes | Yes | Yes | Yes | Yes | Yes | Yes |
|  | *bla*<sub>TEM</sub> gene (plasmid) | No | No | No | No | No | No | No | No | No | No | No | No | No | No |
|  | *tet(M)* (plasmid) | No | No | No | No | No | No | No | No | No | No | No | No | No | No |

<sup>a</sup> Breakpoints according to the European Committee on Antimicrobial Susceptibility testing (www.eucast.org) were used.

## Discussion

In the present study, we show that gonococcal genomes can be *de novo* assembled with high accuracy and contiguity running assemblies with MinION 2D ONT reads in combination with Illumina reads using MaSuRCA. These assemblies varied from one to eight contigs and could be further investigated using tools like Circlator^52^ to obtain circular and finished gonococcal genomes. Furthermore, it is possible to obtain accurate *de novo* assemblies for AMR determinant detection by initially performing Illumina-based correction of individual ONT reads and subsequently using the reads for *de novo* assembly. However, this strategy makes the ONT platform dependent on additional Illumina sequencing. Nevertheless, this is another approach for producing accurate finished genomes and, accordingly, an alternative to PacBio sequencing of microbial reference genomes.

When relying only on the ONT data, the assemblies contained high numbers of mismatches (27-607 mismatches per 100 kb) and indels (641-1125 indels per 100 kb), using even the best ONT assemblies obtained using SMARTdenovo, Canu and Miniasm with 2D ONT reads. There are several possible reasons for the high and incorrect number of CDSs in the ONT assemblies using Prokka including sequencing artefacts, homopolymers and overall high error rate (SNPs, insertions and deletions) in the ONT reads causing the assemblies to contain incorrect internal stop codons and false pseudogenes. Generally, the 2D ONT reads generated more accurate assemblies than the 1D ONT reads and were used in the majority of our downstream analysis. We aimed to obtain highly accurate *de novo* assemblies to be able to rapidly identify relevant gonococcal AMR determinants to predict AMR, using only user-friendly and rapid online sequence analysis tools such as WGSA (www.wgsa.net)^21^ and PubMLST (www.pubmlst.org). Using these online tools, the characterisation of AMR determinants was generally inaccurate using ONT assemblies and, as expected, highly accurate with hybrid assemblies. However, the *in house*-customised CLC Genomics Workbench workflow, including the NG-STAR database^56^, provided 100% concordance with the AMR determinants of the PacBio sequenced 2016 WHO gonococcal reference genomes^41^ using only 2D ONT reads. We can also extract additional AMR determinants and in general genes of interest that are not included in the online tools, because our AMR database is fully customisable. The software workflow is based on extraction of the genes of interest in the customised database from a *de novo* assembly using BLAST algorithms optimised for each AMR determinant and reporting the highest hit, i.e. not only the 100% hit, in also genes with premature stop codons. This is essential because the ONT *de novo* assemblies are prone to errors. The main limitations of any CLC Genomics Workbench are that it is commercial (not an open-source online tool) and it has fairly high system requirements (16-32 GB RAM). Nevertheless, it has a simple general user interface for users who are not familiar with command line bioinformatics and is available to all widely used operating systems. Hopefully, open-source tools and public databases that can handle error-prone assemblies will be inspired by this approach and incorporate e.g. NG-STAR data^56^ rendering costly software obsolete. NG-STAR can currently not use genome assemblies as input for determining the AMR profiles, which limits its possibilities when performing WGS.

Easy and rapid genome sequencing, using platforms such as the ONT MinION, in combination with algorithms that can appropriately predict AMR profiles using only genetic data could be very valuable in future surveillance of gonococcal AMR and spread of AMR gonococcal strains, nationally and internationally. Using the customised CLC Genomics Workbench, we showed that all isolates expressing decreased susceptibility to extended-spectrum cephalosporins (ESCs; cefixime and ceftriaxone) were carrying a mosaic *penA* allele or a *penA* A501 substitution, which are associated with decreased susceptibility or resistance to ESCs. Furthermore, one azithromycin resistant isolate (MIC=4 mg/L) contained the C2611T mutation in four alleles of the 23S rRNA gene, conferring azithromycin resistance. This macrolide-resistance mutation can be challenging to detect accurately in gonococci because of the four alleles of this gene in the gonococcal genome. The number of 23S rRNA gene alleles containing the C2611T mutation is associated with the level of resistance to azithromycin. We included a separate analysis to detect the frequency of the mutated 23S rRNA gene alleles by mapping the 2D ONT reads to a *23S rDNA* reference and analysing the frequency of variation across the gene. We also showed that all ciprofloxacin resistant isolates contained mutations in the *gyrA* gene (resulting in amino acid substitution S91F) and isolates that also had mutations in *parC* (resulting in amino acid substitutions D86N, S87R and E91G) displayed high-level resistance to ciprofloxacin. Finally, the decreased susceptibility and resistance to penicillin could mainly be explained by mutations in *penA* and *ponA,* while resistance to tetracycline appeared to be due to factors other than only *rpsJ* and the *tet(M)* carrying plasmid.

The use 2D ONT reads for phylogenomics of gonococci, e.g. for molecular epidemiological surveillance purposes, would be exceedingly valuable for outbreak investigations and monitoring in nearly real time in local, national and international gonococcal surveillance programmes, which aim to replace traditional, labour intensive and less accurate genotyping techniques such as *N. gonorrhoeae* multi-antigen sequence typing (NG-MAST) and multi-locus sequence typing (MLST) with WGS techniques^21^. Accordingly, we examined the accuracy of performing phylogenomic analysis using 2D ONT reads mapped to a reference genome using BWA-MEM with the nanopore option, and building a phylogeny using the multiple sequence alignment. The ONT data produced a phylogenetic tree topology that was comparable with the one using the Illumina dataset and similarly clustered all related isolates (Fig. 3). However, the number of isolates was limited and the genomic heterogeneity of the strains was high, which might have slightly biased our analysis.

Interestingly, the clustering of isolates was not highly dependent on the number of 2D reads (Fig. 1). This suggests that the read length provides sufficient genome coverage even when a relatively low number of reads are available. For example, the lower number of 2D reads for WHO G (Fig 1) still provided a 99.96-100% genome fraction (Supplementary Table) and, accordingly, the read length compensated for the low number of reads. For the different WHO reference strains, 89.93-100% fraction of the PacBio-sequenced genomes were covered by the ONT reads (Supplementary Table). Consequently, for some purposes including gonococcal phylogenomic analysis, the MinION sequencer has likely collected enough reads in less than one day, which opens up new possibilities in running WGS to investigate gonorrhoea outbreaks in nearly real time, as well as directly from clinical samples.

The main benefits of using the MinION for genome sequencing included the very rapid turn-around time, high accessibility by connecting the small hand-held device to a laptop, and low cost of the sequencer. The main limitations included the high cost of each sequencing run and that the sequencing reads remain less accurate and consistent than Illumina and PacBio sequencing reads.

In conclusion, we show, in the first MinION study for gonococci, that ONT sequences analysed with currently existing open-access, web-based sequence analysis tools are not sufficiently accurate to identify key gonococcal AMR determinants. However, using an appropriate analysis workflow such as an *in house*-developed CLC Genomics Workbench, we show that ONT sequence data can be used for rapid and accurate identification of AMR determinants to predict decreased susceptibility or resistance to recommended therapeutic antimicrobials. We also show that ONT sequence data can be useful for phylogenomics of *N. gonorrhoeae*, e.g. for molecular epidemiological investigations in nearly real time, and, using ONT-Illumina hybrid assemblies, for producing contiguous assemblies and finished gonococcal reference genomes.

## Acknowledgment

We are grateful to Sara Kasraian for laboratory work in RaDAR-Go and Dianne Egli-Gany for her contribution to the management of RaDAR-Go and organisation of the specimens for this study.

We are also grateful for the financial support provided by Örebro County Council Research Committee and the Foundation for Medical Research at Örebro University Hospital, Örebro, Sweden and the SwissTransMed initiative (Translational Research Platforms in Medicine, project number #25/2013: Rapid Diagnosis of Antibiotic Resistance in Gonorrhoea, RaDAR-Go) from the Rectors’ Conference of the Swiss Universities (CRUS). LSB and SRH were supported by Wellcome Grant number 098051. We would like to thank the Pathogen Informatics Group at the Wellcome Sanger Institute for support.

## Authors’ contributions

D.G., V.D., A.E., N.L. and M.U. planned and designed the study. V.D. performed the MinION sequencing. D.G. performed the Illumina sequencing and all the bioinformaticanalysis, with support from S.H. and L.S-B. D.G. supported by M.U wrote a draft of the manuscript. All authors commented on and approved the final manuscript.

